# PET and CSF amyloid-β status are differently predicted by patient features: Information from discordant cases

**DOI:** 10.1101/673467

**Authors:** Juhan Reimand, Arno de Wilde, Charlotte E. Teunissen, Marissa Zwan, Albert D. Windhorst, Ronald Boellaard, Frederik Barkhof, Wiesje M. van der Flier, Philip Scheltens, Bart N.M. van Berckel, Rik Ossenkoppele, Femke Bouwman

## Abstract

**Background:** Amyloid-β PET and CSF Aβ_42_ yield discordant results in 10-20% of patients, possibly providing unique information. Although the predictive power of demographic, clinical, genetic and imaging features for amyloid-positivity has previously been investigated, it is unknown whether these features differentially predict amyloid-β status based on PET or CSF, or whether this differs by disease stage.

**Methods:** We included 768 patients (subjective cognitive decline (SCD, n=194), mild cognitive impairment (MCI, n=127), dementia (AD and non-AD, n=447) with amyloid-β PET and CSF Aβ_42_ measurement within one year. 97(13%) patients had discordant PET/CSF amyloid-β status. We performed parallel random forest models predicting separately PET and CSF status using 17 patient features (demographics, APOE4 positivity, CSF (p)tau, cognitive performance, and MRI visual ratings) in the total patient group and stratified by syndrome diagnosis. Thereafter, we selected features with the highest variable importance measure (VIM) as input for logistic regression models, where amyloid status on either PET or CSF was predicted by (i) the selected patient feature, and (ii) the patient feature adjusted for the status of the other amyloid modality.

**Results:** APOE4, CSF tau and p-tau had highest VIM for PET and CSF in all groups. In the amyloid-adjusted logistic regression models, p-tau was a significant predictor for PET-amyloid in SCD (OR=1.02[1.01-1.04], p_FDR_=0.03), MCI (OR=1.05[1.02-1.07], p_FDR_<0.01) and dementia (OR=1.04[1.03-1.05], p_FDR_<0.001), but not for CSF-amyloid. APOE4 (OR=3.07[1.33-7.07], p_unc_<0.01) was associated with CSF-amyloid in SCD, while it was only predictive for PET-amyloid in MCI (OR=9.44[2.93,30.39], p_FDR_<0.01). Worse MMSE scores (OR=1.21[1.03-1.41], p_unc_=0.02) were associated to CSF-amyloid status in SCD, whereas worse memory (OR=1.17[1.05-1.31], p_FDR_=0.02) only predicted PET positivity in dementia.

**Conclusion:** Amyloid status based on either PET or CSF was predicted by different patient features and this varied by disease stage, suggesting that PET-CSF discordance yields unique information. The stronger associations of both APOE4 carriership and worse memory z-scores with CSF-amyloid in SCD suggests that CSF-amyloid is more sensitive early in the disease course. The higher predictive value of CSF p-tau for a positive PET scan suggests that PET is more specific to AD pathology. These findings can influence the choice between amyloid biomarkers in future studies or trials.

## INTRODUCTION

Alzheimer’s disease (AD) is characterized by the accumulation of amyloid-β plaques, which has been shown to occur decades before symptom onset (1,2). Amyloid-β pathology can be detected *in vivo* by positron emission tomography (PET) using amyloid-β radiotracers such as [^11^C]Pittsburgh compound-B (PIB), [^18^F]Florbetapir, [^18^F]Florbetaben or [^18^F]Flutemetamol allows to directly visualize fibrillary amyloid-β deposits in brain tissue (3–6). Alternatively, Aβ_42_ levels in cerebrospinal fluid (CSF) reflect the concentration of soluble amyloid-β, which correlates with cerebral amyloid-β depositions (7). PET and CSF have been included as equal alternatives into diagnostic criteria for both research (2,8,9) and clinical practice (10–12). However, it has been repeatedly shown that in 10-20% of patients these modalities yield conflicting results (13–15). This discordance may include valuable information on underlying clinical or neuropathological differences (16).

A combination of various patient features has previously been demonstrated to predict amyloid-β positivity based on PET and/or CSF (17,18). In particular, a combination of demographic information, *APOE* ε4 carriership, neuropsychological tests, and magnetic resonance imaging (MRI) measures was effective in predicting amyloid-β status (19). Additionally, CSF tau and p-tau have been shown to be predictive of amyloid PET status (20). So far it has not been investigated whether the predictive ability of patient features for amyloid-β pathology differs when detected by PET or by CSF. If these two modalities would provide partially independent information about the underlying pathology, then this could be reflected by the differences in predictive patterns for the two modalities. Additionally, as it has been suggested that CSF might be able to detect amyloid-β depositions earlier (21), it is possible that the relative predictive contribution of a patient feature changes throughout the course of Alzheimer’s disease. Therefore, we investigate the unique information provided by the PET-CSF discordant population using the predictive patterns for amyloid PET and CSF in (i) the total patient group and (ii) stratifying by syndrome diagnosis. Exploring this allows us to gain insight in the clinical and neurobiological factors related to discordant results between amyloid-β PET and CSF and ultimately about the underlying neuropathological processes during the disease course of AD.

## METHODS

### Study Population

We retrospectively included 777 patients, who had visited our tertiary memory clinic between 2005 and 2017 and had undergone both CSF Aβ_42_ analysis and amyloid-β PET within one year. We excluded nine patients that did not pass PET imaging quality control. Patients were screened according to the standardized protocol of the Amsterdam Dementia Cohort (22,23). This includes a clinical and neuropsychological evaluation, *APOE* genotyping, MR imaging and a lumbar puncture for CSF analysis. Patient diagnosis was determined during a multidisciplinary meeting, according to international guidelines (10,11,24–32).

### Neuropsychological testing

Subjects underwent extensive neuropsychological testing as part of their diagnostic process. Mini-Mental State Examination (MMSE) scores were used to measure global cognition. In addition, five cognitive domains were assessed (33). We used the visual association test (VAT), total immediate recall, and the Dutch version of the Rey Auditory Verbal Learning test (delayed recall) to assess memory. Language was assessed by VAT naming and category fluency (animals). The Trail-Making Test (TMT) part A, Digit Span forwards and the Stroop test I and II were used for attention. Executive functioning was assessed by TMT B, Digit Span backwards, Stroop test III, the Frontal Assessment Battery, and the Dutch version of the Controlled Oral Word Association Test (letter fluency). Finally, we assessed visuospatial functioning by Visual Object and Space Perception battery: tests incomplete letters, dot counting and number location.

For every test, we derived Z-scores using the mean and standard deviation values from a group of healthy controls (n = 360) (33). TMT-A, TMT-B and Stroop Test scores were log-transformed to account for the non-normal distribution of the data and multiplied by −1 so that lower scores would indicate worse performance. In case TMT B was aborted and TMT A was available (n = 132), we estimated the TMT B score using the multiplication of TMT A score with mean TMT B/A score ratio from the respective diagnostic group (34). Thereafter, based on available tests we used z-scores to compile a composite score for each of the five cognitive domains.

### CSF

CSF was obtained by lumbar puncture between L3/4, L4/5 or L5/S1 intervertebral space, using a 25-gauge needle and a syringe (35). The samples were collected in polypropylene microtubes and centrifuged at 1800g for 10min at 4°C. Thereafter, the samples were frozen at −20 °C until manual analyses of Ab_42_, tau and p-tau were performed using sandwich ELISAs [Innotest assays: β-amyloid1-42, tTAU-Ag and PhosphoTAU-181p; Fujirebio (formerly Innogenetics)] at the Neurochemistry Laboratory of the Department of Clinical Chemistry of VUmc. As the median CSF Aβ_42_ values of our cohort have been gradually increasing over the years (36), we determined CSF amyloid-β status using Aβ_42_ values that had been adjusted for the longitudinal upward drift. We used a uniform cut-off of 813 pg/mL to dichotomize CSF data, as continuous measures were not available for PET imaging (37).

### PET

Amyloid-β PET scanning is not part of standard diagnostic process in the Amsterdam Dementia Cohort. Patients underwent an amyloid-β PET for research purposes in the vast majority (38–43) or otherwise in case of a diagnostic dilemma. Amyloid-β PET scans were performed using the following PET scanners: ECAT EXACT HR+ scanner (Siemens Healthcare, Germany) and Gemini TF PET/CT, Ingenuity TF PET-CT and Ingenuity PET/MRI (Philips Medical Systems, the Netherlands). We included PET scans using four different radiotracers: [^18^F]Florbetaben (38,43) (n=322, 42%), [^11^C]PIB (40–42) (n=271, 35%), [^18^F]Flutemetamol (44) (n=151, 20%), and [^18^F]Florbetapir (39) (n=24, 3%). PET scans were rated as positive or negative based on visual read by an expert nuclear medicine physician (BvB). PET scans were performed, on average, within 54 (±75) days of the lumbar puncture.

### MRI

The acquisition of MRI scans has been extensively described previously (23). During the period of 2005 to 2017, the following scanners have been used: Discovery MR750 and Signa HDXT (both GE Medical Systems, USA); Ingenuity TF PET/MR (Philips Medical Systems, The Netherlands); Titan (Toshiba Medical Systems, Japan); Magnetom Impact and Sonata (Siemens Healthcare, Germany). The MRI protocol included 3D T1-weighted, T2-weighted, fluid-attenuated inversion recovery (FLAIR), gradient-echo T2* and/or susceptibility weighted imaging sequences. The scans were visually assessed by a neuroradiologist on three different image planes. Parietal atrophy was rated using the posterior cortical atrophy (PCA) scale (45), medial temporal atrophy using the medial temporal lobe atrophy (MTA) scale (46) and the extent of white matter hyperintensities according to the Fazekas scale (47). MTA and PCA scores were scored separately for right and left and averaged thereafter. In addition, the scans were assessed for the existence of lacunes and microbleeds.

### Patient groups

We stratified the patients based on syndrome diagnosis: subjective cognitive decline (SCD, n=194 (29%)) (48), mild cognitive impairment (MCI, n=127 (17%)), and dementia (n=447 (58%)). Within the dementia group, 309 (69%) patients had the diagnosis of Alzheimer’s disease, 66 (15%) a diagnosis within the frontotemporal dementia spectrum, 22 (5%) dementia with Lewy bodies, 6 (1%) vascular dementia and 44 (10%) other dementia syndromes. To reflect the information provided to the models in our analysis, we present patient group characteristics based on the binarized amyloid-β status on PET and CSF: concordantly positive (PET+/CSF+) or negative (PET-/CSF-for amyloid-β pathology, or discordantly positive amyloid-β status based on PET (PET+/CSF-) or CSF (PET/CSF+).

### Statistical Analysis

Statistical analysis was performed using R software (Version 3.4.4) (49). When presenting our study population by binarized PET/CSF status groups, we compared patient features using Chi-squared tests, two samples *t*-tests, Wilcoxon Rank-Sum tests and linear regression models with Bonferroni correction for group-wise testing. Cognitive scores were compared while adjusting for age, sex, education and syndrome diagnosis.

All subsequent analyses were performed in the total patient group as well as in the syndrome diagnosis groups of SCD, MCI and dementia. We first summarized the relative predictive power of every variable in predicting PET and CSF amyloid-β status using random forest modelling. As classifier models are affected by missing data, we accounted for missing values using multiple imputations (with 25 imputations and 5 iterations) (**Supplementary Table 1**). For each of the imputed dataset, we ran two conditional random forest models (50,51), predicting separately PET and CSF status using various patient features associated with Alzheimer’s disease (17–19). As predictors, we selected demographic information (age, sex, education), biomarkers (*APOE* ε4 positivity, CSF tau and p-tau), cognitive measures (MMSE; z-scores for memory, language, attention, executive, visuospatial), and MRI scores (MTA, PCA, Fazekas scale, the presence of lacunes and microbleeds). We used the area-under-the-curve (AUC)-based permutation variable importance measure (VIM) to estimate the relative predictive power for every patient feature.

This measure was selected because of its higher accuracy in datasets with an unbalanced outcome class (52) and we expected this to be especially helpful in the SCD group with a low prevalence of amyloid-β positivity. Accuracy, sensitivity and specificity of the random forest models were evaluated using the mean out-of-bag error estimates. Using this method, the performance of every tree in the random forest model is evaluated on the approximately 37% of observations that are not used for its training (53).

For the second stage of the analysis, we selected patient features based on their predictive value in the random forest models. Similar to a previous study (19), we included patient features when their median VIM over the 25 random forests models for predicting either PET or CSF was higher than the median VIM of all the features for the patient group. First, using two sample t-testing of 1000x bootstrapped samples with replacement, we compared the VIM of every selected patient feature between the parallel random forest models predicting amyloid-β PET and CSF status. Secondly, to determine the unadjusted predictive power of these patient features, we performed bivariate logistic regression models with either PET or CSF positivity as the outcome and the selected patient features as predictors. Thirdly, to investigate the added predictive value of a patient feature to the other amyloid-β modality, we performed multivariable logistic regression models, with either PET or CSF positivity as the outcome and the selected patient feature with the status of the other amyloid-β modality as predictors.

We calculated the odds ratios (OR) with corresponding 95% confidence intervals for every patient feature both in the original dataset and in the 25x imputed datasets. Non-overlapping confidence intervals were considered significantly different. We used the False discovery rate (FDR) correction with a significance level of 0.05 to account for multiple testing (54).

## RESULTS

### Overview of features

Patient characteristics grouped by PET/CSF status are summarized in **Table 1** and CSF Aβ_42_ levels shown in **Figure 1**. 32 patients (4%) were discordantly amyloid-β positive based on PET and 65 (8%) based on CSF. In general, the PET+CSF+ group showed a higher proportion of APOE ε4 carriers, more AD-like CSF markers, MRI features, and lower cognitive scores compared to PET-CSF-group. CSF tau and p-tau were lower in both PET-CSF- and PET-CSF+ groups, compared to PET+CSF- and PET+CSF+. The PET-CSF-group contained a lower proportion of *APOE* ε4 carriers and better cognitive scores than patients in the discordant groups.

**Table 1.**
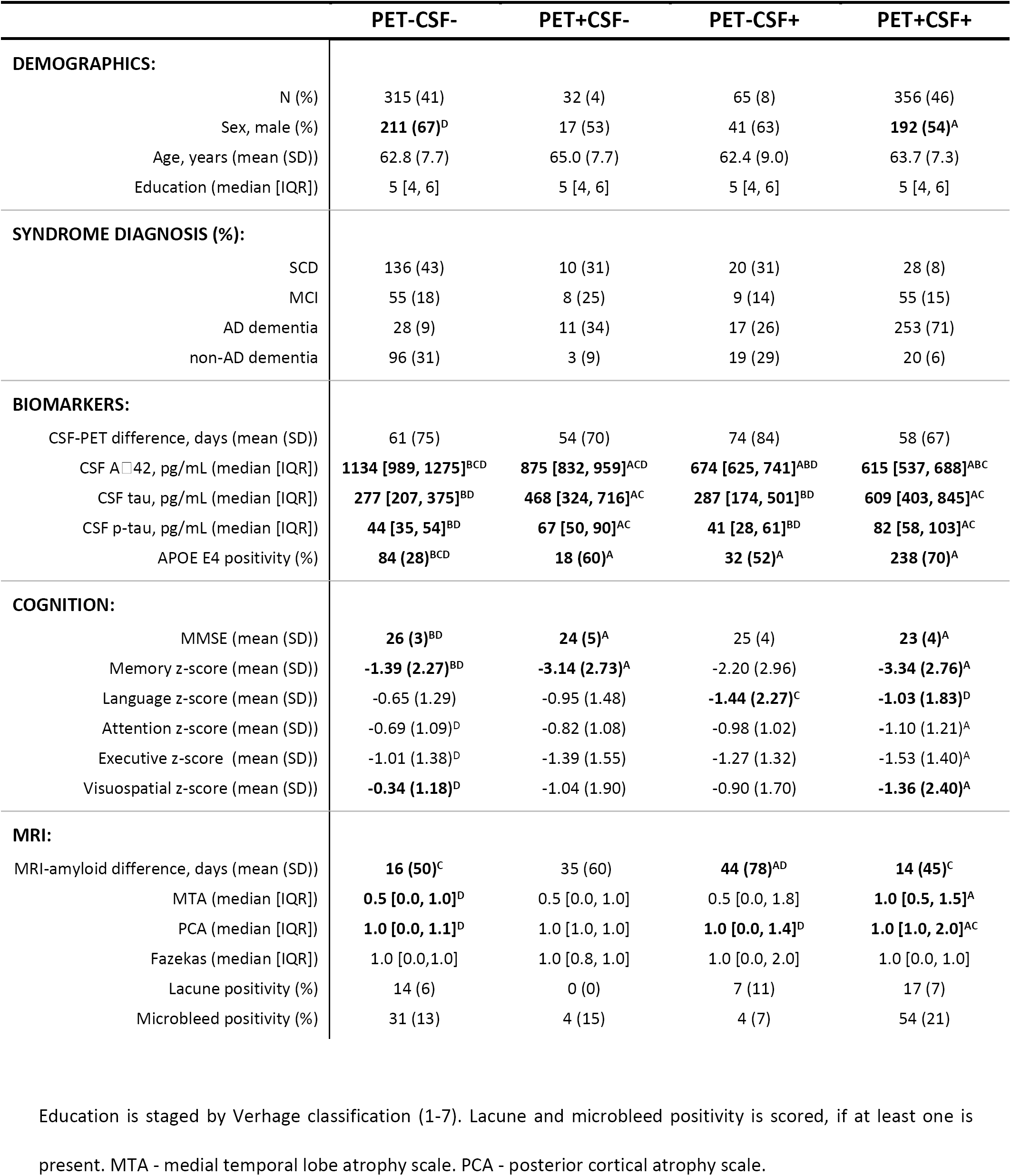
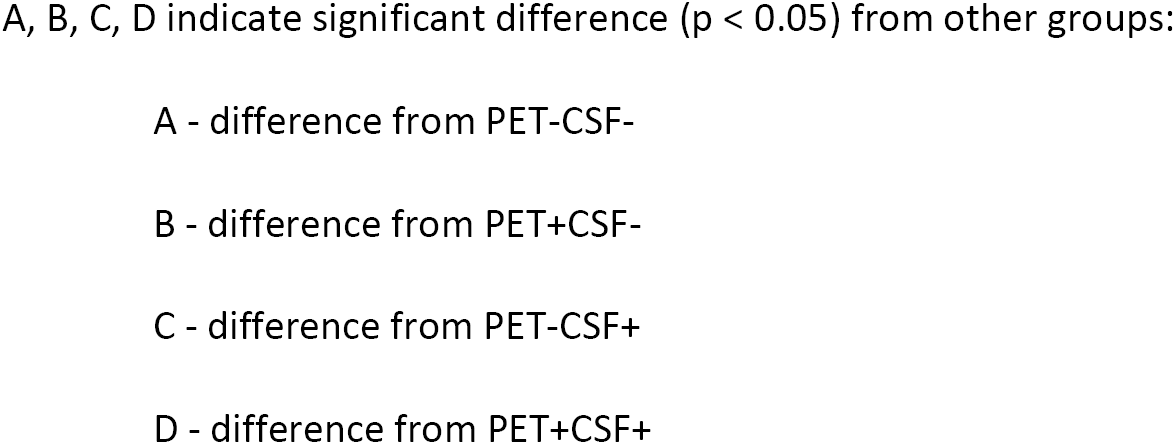
Patient groups by PET/CSF amyloid status.

**Figure 1.**
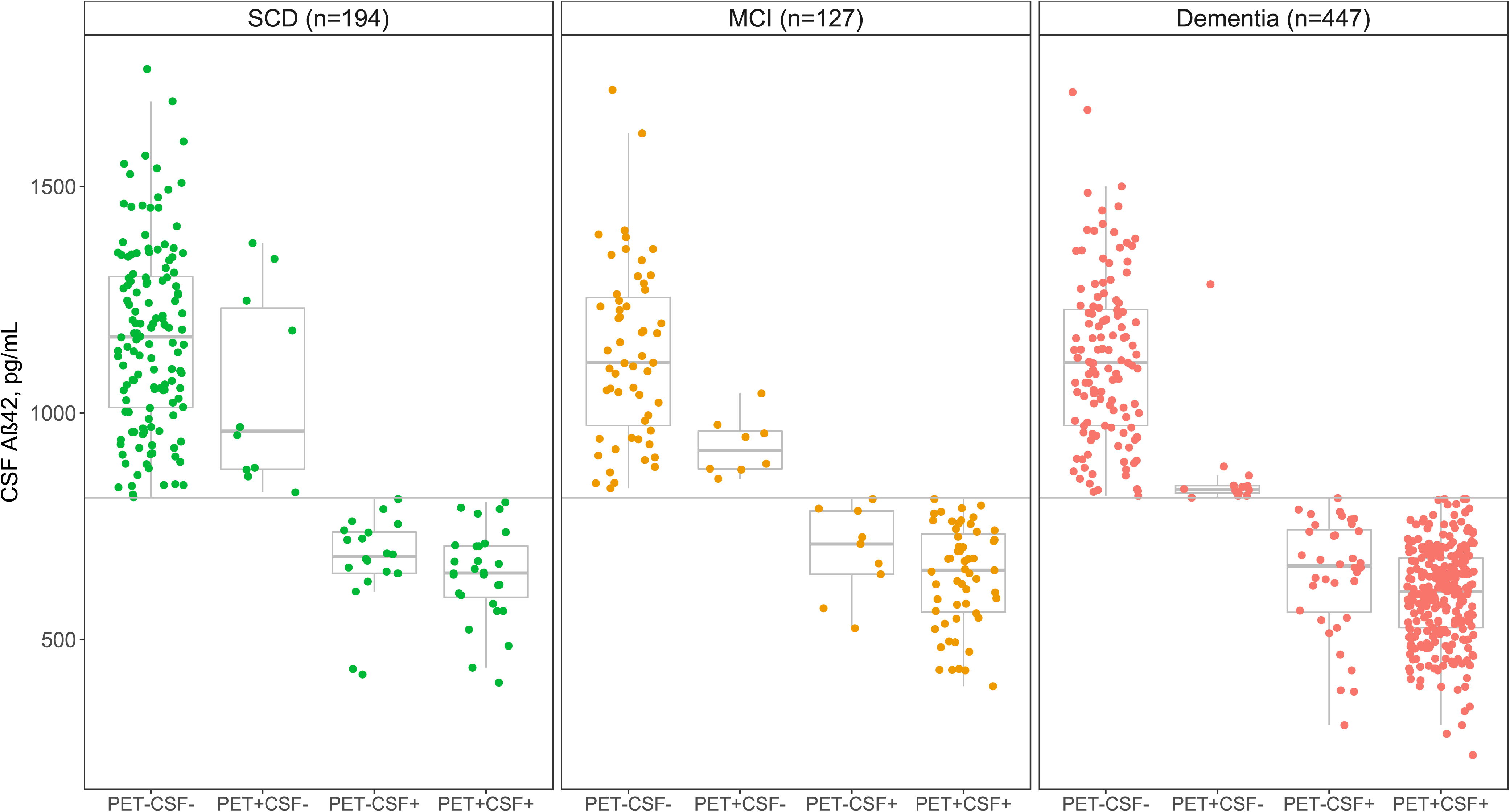
CSF Aβ_42_ values by PET/CSF amyloid status groups in SCD, MCI and dementia. Horizontal line indicates the cut-off of 813 pg/mL used for dichotomization of CSF-amyloid.

### Patient feature selection

We performed random forest modelling to (i) select patient features based on overall VIM for multivariable logistic regression models and (ii) compare the VIM between models predicting PET and CSF amyloid-β status. Summarized VIM values over the 25 random forest models (one with each set of imputed data) are shown in **Figure 2**. Out-of-bag accuracy, sensitivity and specificity rates for the random forest models are reported in **Supplementary Table 2.** *APOE* ε4 positivity was the most important predictor for amyloid-β positivity in the total patient group for both PET (VIM=0.28±0.14 (mean VIM*100 ± standard deviation)) and CSF (VIM=0.31±0.19, PET/CSF VIM difference: p=0.51). CSF tau was similarly important when predicting PET (VIM=0.06±0.03) or CSF (VIM=0.06±0.03, p=0.51), but CSF p-tau was a more important predictor for PET (VIM=0.12±0.03) compared to CSF (VIM=0.03±0.01, p<0.001).

**Figure 2.**
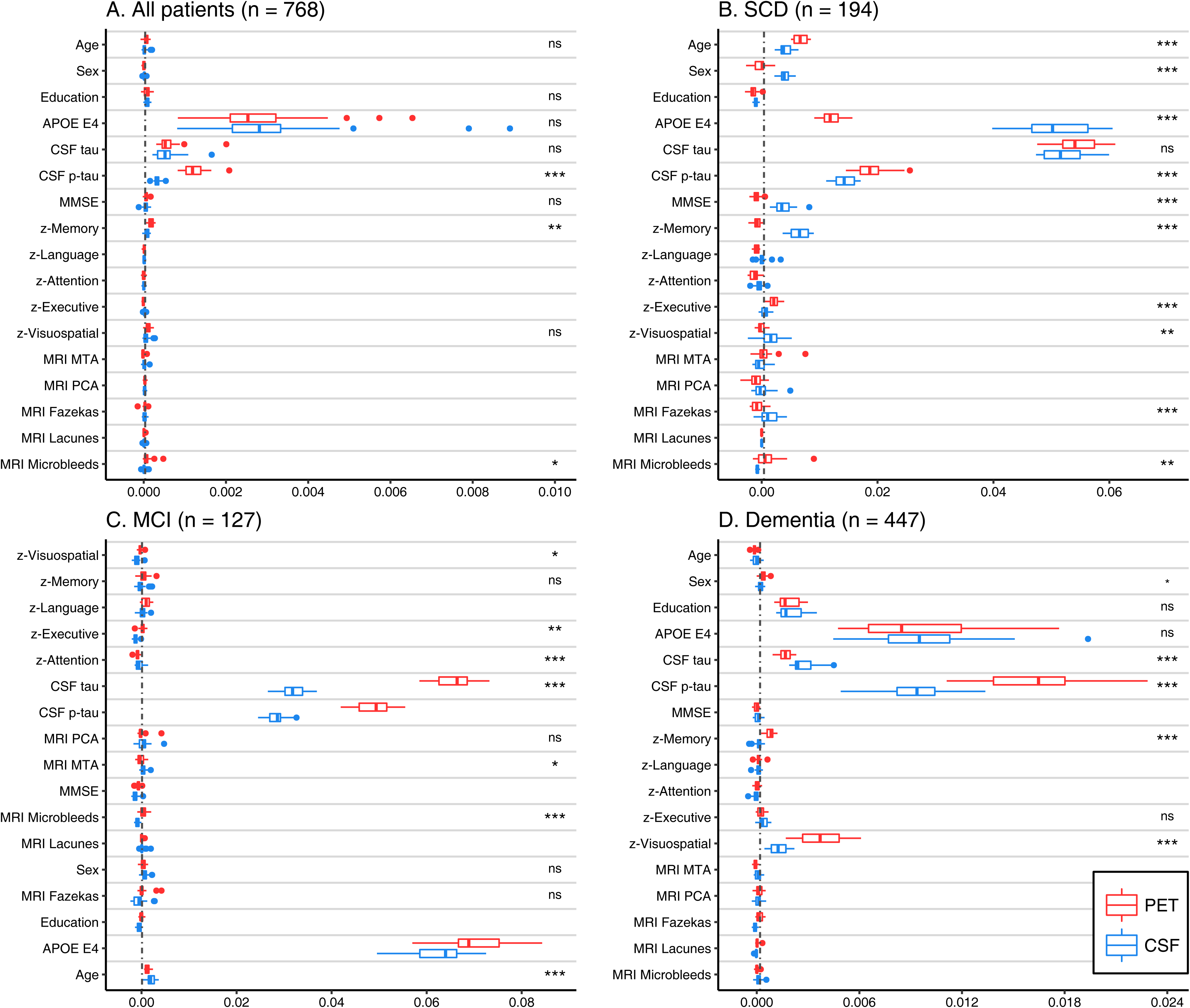
Relative predictive power of patient features for amyloid PET and CSF status. AUC-based variable importance (VIM) from 25 random forest models predicting PET status and 25 models from predicting CSF status are plotted. P-values (*** - p<0.001, ** - p<0.01, * - p<0.05, ns - non-significant) indicate the bootstrapped difference of VIM values between models predicting PET and CSF status.

Subsequently, we stratified for syndrome diagnosis. In SCD, *APOE* ε4 positivity was a stronger predictor for CSF (VIM=5.06±0.63) than PET (VIM=1.19±0.18, p<0.001), whereas CSF p-tau was more associated with PET (VIM=1.89±0.29) than CSF amyloid-β status. (VIM=1.44±0.18, p<0.001). Additionally, MMSE (VIM=0.37±0.17) and memory score (VIM=0.66±0.18) had a stronger association with CSF than PET (both p<0.001). CSF tau was equally important for predicting PET (VIM=5.47±0.36) or CSF amyloid-β status (VIM=5.24±0.38, p=0.13). In contrast to the findings in SCD, in MCI, A*POE* ε4 carriership was a stronger predictor for PET (VIM=7.07±0.69) than for CSF (VIM=6.31±0.56, p<0.01). Moreover, CSF tau and p-tau were more important for predicting PET (respectively VIM=6.56±0.42; and VIM=4.90±0.38) than for CSF amyloid-β status (VIM=3.20±0.25, p<0.001; and VIM=2.84±0.21, p<0.001). In dementia, CSF p-tau was more predictive of PET (VIM=1.60±0.32) than CSF (VIM=0.92±0.22, p<0.001), but CSF tau was a stronger predictor for CSF (VIM=0.27±0.07) than for PET amyloid-β status (VIM=0.17±0.04, p<0.001). Both PET (VIM=0.93±0.34) and CSF (VIM=0.97±0.35, p=0.49) had a strong association to *APOE* ε4 carriership. Finally, visuospatial (VIM=0.37±0.13) and memory (VIM=0.08±0.03) scores were more important for predicting PET positivity (both p<0.001).

We verified the predictive ability of the selected patient features with bivariate logistic regression models for PET and CSF status (**Table 2;** all possible models in **Supplementary Table 3**). The bivariate models largely confirmed the feature selection of the random forest procedure, as *APOE* ε4, CSF tau and CSF p-tau were consistently significant predictors in all groups. In the total group and dementia, most of the patient features selected based on the random forest models were significant predictors.

**Table 2.** Predictive value of patient features for amyloid status based on PET or CSF.

|  |  | TOTAL |  |  |  |  | SCD |  |  |  |  | MCI |  |  |  |  | DEMENTIA |  |  |  |  |  |  |  |
| --- | --- | --- | --- | --- | --- | --- | --- | --- | --- | --- | --- | --- | --- | --- | --- | --- | --- | --- | --- | --- | --- | --- | --- | --- |
| Predictor | Out-<br>come | Odds ratio<br>(95% CI) | p<br>unc | p<br>FDR | Imputed |  | Odds ratio<br>(95% CI) | p<br>unc | p<br>FDR | Imputed |  | Odds ratio<br>(95% CI) | p<br>unc | p<br>FDR | Imputed |  | Odds ratio<br>(95% CI) | p<br>unc | p<br>FDR | Imputed |  | Odds ratio<br>(95% CI) | p<br>unc | p<br>FDR |
|  |  |  |  |  | Odds ratio<br>(95% CI) | p<br>FDR |  |  |  | Odds ratio<br>(95% CI) | p<br>FDR |  |  |  | Odds ratio<br>(95% CI) | p<br>FDR |  |  |  | Odds ratio<br>(95% CI) | p<br>FDR |  |  |  |
| Age | PET | 1.02(1.00,1.04) |  |  | 1.02(1.00,1.04) |  | 1.06(1.01,1.12) | * |  | 1.06(1.01,1.12) |  | 0.96(0.92,1.00) |  |  | 0.96(0.92,1.00) |  |  |  |  |  |  |  |  |  |
|  | CSF | 1.01(0.99,1.03) |  |  | 1.01(0.99,1.03) |  | 1.04(1.00,1.09) |  |  | 1.04(1.00,1.09) |  | 0.96(0.92,1.00) |  |  | 0.96(0.92,1.00) |  |  |  |  |  |  |  |  |  |
| Sex, F | PET |  |  |  |  |  | 1.64(0.81,3.36) |  |  | 1.64(0.81,3.36) |  | 2.45(1.14,5.26) | * |  | 2.45(1.14,5.26) |  | 1.70(1.13,2.53) | ** | * | 1.70(1.13,2.53) | * |  |  |  |
|  | CSF |  |  |  |  |  | 1.91(0.99,3.69) |  |  | 1.91(0.99,3.69) |  | 2.01(0.94,4.28) |  |  | 2.01(0.94,4.28) |  | 1.40(0.93,2.12) |  |  | 1.40(0.93,2.12) |  |  |  |  |
| Education | PET | 1.06(0.94,1.19) |  |  | 1.07(0.95,1.20) |  |  |  |  |  |  |  |  |  |  |  | 1.28(1.08,1.52) | ** | ** | 1.30(1.10,1.53) | ** |  |  |  |
|  | CSF | 1.04(0.93,1.17) |  |  | 1.05(0.94,1.18) |  |  |  |  |  |  |  |  |  |  |  | 1.29(1.08,1.54) | ** | ** | 1.30(1.09,1.54) | ** |  |  |  |
| APOE E4 | PET | 4.72(3.46,6.44) | *** | *** | 4.57(3.34,6.24) | *** | 2.97(1.42,6.20) | ** | * | 2.97(1.42,6.19) | * | 14.55(6.08,34.82) | *** | *** | 13.43(5.62,32.11) | *** | 3.63(2.39,5.50) | *** | *** | 3.51(2.30,5.34) | *** |  |  |  |
|  | CSF | 4.60(3.36,6.28) | *** | *** | 4.46(3.26,6.12) | *** | 3.82(1.90,7.70) | *** | *** | 3.75(1.86,7.57) | ** | 8.28(3.70,18.54) | *** | *** | 7.76(3.49,17.27) | *** | 3.68(2.39,5.69) | *** | *** | 3.59(2.33,5.53) | *** |  |  |  |
| CSF tau | PET | 1.005<br>(1.004,1.006) | *** | *** | 1.005<br>(1.004,1.006) | *** | 1.004<br>(1.002,1.006) | *** | *** | 1.004<br>(1.002,1.006) | *** | 1.008<br>(1.005,1.011) | *** | *** | 1.008<br>(1.005,1.011) | *** | 1.004<br>(1.003,1.005) | *** | *** | 1.004<br>(1.003,1.005) | *** |  |  |  |
|  | CSF | 1.004<br>(1.003,1.005) | *** | *** | 1.004<br>(1.003,1.005) | *** | 1.003<br>(1.002,1.005) | *** | *** | 1.003<br>(1.002,1.005) | ** | 1.003<br>(1.002,1.005) | *** | *** | 1.003<br>(1.002,1.005) | *** | 1.004<br>(1.003,1.004) | *** | *** | 1.003<br>(1.002,1.004) | *** |  |  |  |
| CSF p-tau | PET | 1.05(1.04,1.06) | *** | *** | 1.05(1.04,1.06) | *** | 1.04(1.02,1.05) | *** | *** | 1.04(1.02,1.05) | *** | 1.05(1.03,1.07) | *** | *** | 1.05(1.03,1.07) | *** | 1.05(1.04,1.06) | *** | *** | 1.05(1.04,1.06) | *** |  |  |  |
|  | CSF | 1.04(1.03,1.04) | *** | *** | 1.04(1.03,1.04) | *** | 1.03(1.01,1.04) | *** | *** | 1.02(1.01,1.04) | ** | 1.03(1.01,1.04) | *** | *** | 1.03(1.01,1.04) | *** | 1.04(1.03,1.05) | *** | *** | 1.04(1.03,1.05) | *** |  |  |  |
| MMSE | PET | 1.20(1.15,1.25) | *** | *** | 1.19(1.14,1.24) | *** | 1.03(0.89,1.19) |  |  | 1.02(0.88,1.18) |  |  |  |  |  |  |  |  |  |  |  |  |  |  |
|  | CSF | 1.20(1.15,1.25) | *** | *** | 1.19(1.14,1.25) | *** | 1.15(1.01,1.31) | * |  | 1.13(1.00,1.29) |  |  |  |  |  |  |  |  |  |  |  |  |  |  |
| Memory | PET | 1.36(1.27,1.47) | *** | *** | 1.36(1.27,1.46) | *** | 1.13(0.82,1.55) |  |  | 1.14(0.83,1.56) |  | 1.26(1.02,1.57) | * |  | 1.25(1.00,1.54) |  | 1.20(1.10,1.30) | *** | *** | 1.20(1.10,1.31) | *** |  |  |  |
|  | CSF | 1.32(1.23,1.42) | *** | *** | 1.32(1.23,1.42) | *** | 1.23(0.92,1.64) |  |  | 1.22(0.91,1.62) |  | 1.16(0.95,1.42) |  |  | 1.12(0.92,1.38) |  | 1.14(1.05,1.24) | ** | ** | 1.15(1.06,1.26) | ** |  |  |  |
| Language | PET |  |  |  |  |  |  |  |  |  |  | 0.38(0.18,0.81) | * | * | 0.44(0.21,0.95) |  |  |  |  |  |  |  |  |  |
|  | CSF |  |  |  |  |  |  |  |  |  |  | 0.71(0.39,1.27) |  |  | 0.71(0.39,1.29) |  |  |  |  |  |  |  |  |  |
| Executive | PET |  |  |  |  |  | 1.02(0.72,1.45) |  |  | 1.02(0.72,1.44) |  | 0.68(0.45,1.04) |  |  | 0.70(0.46,1.05) |  | 0.95(0.82,1.11) |  |  | 0.96(0.82,1.11) |  |  |  |  |
|  | CSF |  |  |  |  |  | 1.05(0.76,1.44) |  |  | 1.05(0.76,1.44) |  | 0.82(0.55,1.22) |  |  | 0.83(0.55,1.24) |  | 0.88(0.76,1.03) |  |  | 0.89(0.76,1.04) |  |  |  |  |
| Visuo-<br>spatial | PET | 1.36(1.22,1.52) | *** | *** | 1.33(1.19,1.48) | *** | 0.92(0.59,1.45) |  |  | 0.89(0.56,1.41) |  |  |  |  |  |  | 1.30(1.15,1.49) | *** | *** | 1.25(1.10,1.43) | ** |  |  |  |
|  | CSF | 1.38(1.23,1.55) | *** | *** | 1.34(1.19,1.50) | *** | 1.34(0.93,1.93) |  |  | 1.35(0.95,1.93) |  |  |  |  |  |  | 1.21(1.07,1.37) | ** | ** | 1.16(1.03,1.31) | * |  |  |  |
| MRI MTA | PET |  |  |  |  |  |  |  |  |  |  | 0.77(0.48,1.24) |  |  | 0.75(0.47,1.20) |  |  |  |  |  |  |  |  |  |
|  | CSF |  |  |  |  |  |  |  |  |  |  | 1.09(0.69,1.73) |  |  | 0.98(0.62,1.54) |  |  |  |  |  |  |  |  |  |
| MRI PCA | PET |  |  |  |  |  |  |  |  |  |  | 0.84(0.47,1.51) |  |  | 0.85(0.48,1.51) |  |  |  |  |  |  |  |  |  |
|  | CSF |  |  |  |  |  |  |  |  |  |  | 0.74(0.41,1.32) |  |  | 0.75(0.42,1.32) |  |  |  |  |  |  |  |  |  |
| MRI Fazekas | PET |  |  |  |  |  | 0.98(0.53,1.83) |  |  | 0.92(0.50,1.71) |  |  |  |  |  |  |  |  |  |  |  |  |  |  |
|  | CSF |  |  |  |  |  | 1.38(0.79,2.41) |  |  | 1.25(0.72,2.16) |  |  |  |  |  |  |  |  |  |  |  |  |  |  |
| MRI<br>microbleeds | PET | 1.91(1.21,3.01) | ** | ** | 1.59(1.01,2.50) |  | 2.17(0.75,6.30) |  |  | 1.84(0.68,4.99) |  | 1.32(0.52,3.32) |  |  | 1.11(0.47,2.62) |  |  |  |  |  |  |  |  |  |
|  | CSF | 1.51(0.96,2.39) |  |  | 1.35(0.86,2.10) |  | 1.49(0.52,4.25) |  |  | 1.32(0.49,3.52) |  | 1.06(0.42,2.65) |  |  | 0.98(0.41,2.32) |  |  |  |  |  |  |  |  |  |

\*\*\* - p<0.001, \*\* - p<0.01, \* - p<0.05. P-values indicate the significance of the patient feature in the model. Uncorrected p-values and corrected p-values are reported per model, additionally corrected p-values for imputed data. False discovery rate (FDR) correction was performed for multiple comparisons. Cognitive scores have been multiplied by -1, therefore lower scores usually indicate higher odds ratios for amyloid positivity.

### Amyloid-adjusted multivariable logistic regression models

We investigated the added predictive value of the selected patient features to the other amyloid-β modality with multivariable logistic regression models (**Table 3**; all possible models in **Supplementary Table 4**). We assumed that if PET and CSF would truly provide equal information about amyloid status, additional patient features should never be significant predictors in these models, as the other amyloid status would already provide sufficient predictive power. However, if a patient feature added significant information, this would show a stronger association between the feature and the predicted amyloid-β modality.

**Table 3.**
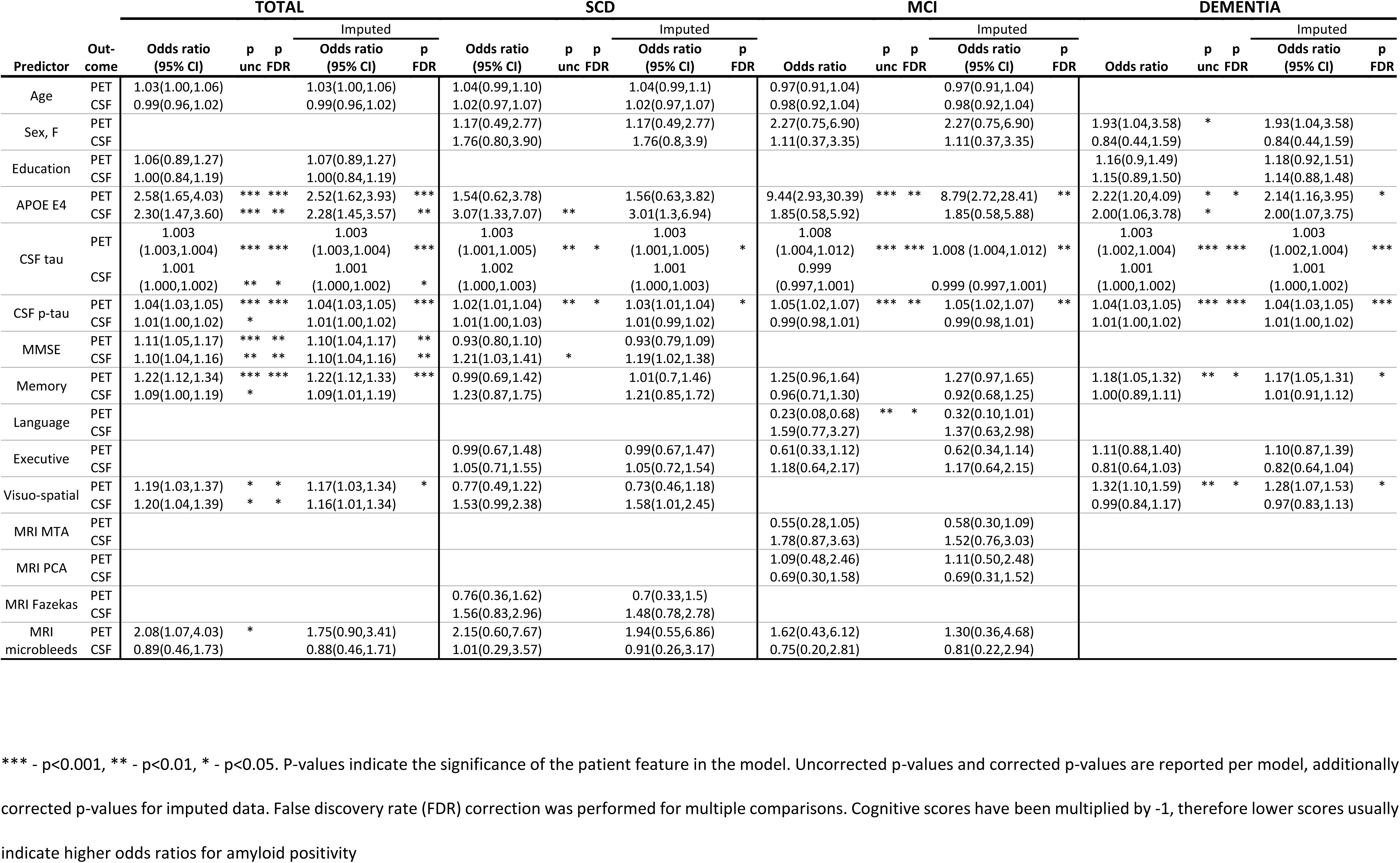
Amyloid-adjusted predictive value of patient features for amyloid status based on PET or CSF.

The following odds ratios are adjusted for the status of the other amyloid modality. In the total group increased levels of CSF p-tau (odds ratio (OR)=1.04[95% confidence interval:1.03,1.05] vs OR=1.01[1.00,1.02]) and tau (OR=1.003[1.003,1.004] vs OR=1.001[1.000,1.002]) were more strongly associated with PET than CSF. In SCD, increased levels of CSF p-tau (OR=1.02[1.01,1.04], p=0.03) and tau (OR=1.003[1.001,1.005], p=0.03) were predictive of only PET, but not CSF positivity (respectively, OR=1.01[1.00,1.03], p=0.42 and OR=1.002[1.000,1.003], p=0.37). APOE ε4 carriership (OR=3.07[1.33,7.07], p_unc_<0.01, p_FDR_=0.07) and lower MMSE scores (OR=1.21[1.03,1.41], p_unc_=0.02, p_FDR_=0.11) showed a predictive trend towards amyloid-β status based on CSF, but not on PET (respectively, OR=1.54[0.62,3.78], p=0.56 and vs OR=0.93[0.80,1.10], p=0.61). In MCI, a positive PET scan was more strongly predicted by *APOE* ε4 carriership (OR=9.44[2.93,30.39], p<0.01 vs OR=1.85(0.58,5.92), p=0.60 for CSF status), and by increased levels of CSF p-tau (OR=1.05[1.02,1.07], p<0.01 vs 0.99[0.98,1.01], p=0.70) and tau (OR=1.008[1.004,1.012], p<0.001 vs OR=0.999[0.997,1.001], p=0.65). Finally, in dementia, PET status had a stronger association with increased levels of CSF p-tau (OR=1.04[1.03,1.05], p<0.001 vs OR=1.01[1.00,1.03], p=0.19 for CSF), tau (OR=1.003[1.002,1.004], p<0.001 vs OR=1.001[1.000,1.002], p=0.13), and with a worse performance in memory (OR=1.18[1.05,1.32], p=0.02 vs OR=1.00[0.89,1.11], p=0.95), and visuospatial ability (OR=1.32[1.10,1.59], p=0.02 vs OR=0.99[0.84,1.17], p=0.95) than CSF amyloid-β status. *APOE* ε4 carriership was similarly associated with both PET (OR=2.22[1.20,4.09], p=0.03) and CSF (OR=2.00[1.06,3.78], p_unc_=0.03, p_FDR_=0.09). No patient feature showed a higher association with CSF in dementia.

## DISCUSSION

We investigated the predictive patterns of various patient features for amyloid-β status based on PET or CSF to determine (i) whether these features have a different association with PET or CSF and (ii) whether this differs per disease stage. We found significant differences in the predictive strength of patient features for amyloid-β status based on PET or CSF. For example, CSF tau, and, especially, CSF p-tau consistently showed a stronger association with amyloid-β status on PET. Additionally, the differential predictive pattern was influenced by the extent of cognitive impairment, as CSF tau was more important in SCD and MCI, while CSF p-tau became more important in the stage of dementia. Moreover, *APOE* ε4 carriership was more predictive towards CSF status in SCD, whereas it was more predictive towards PET in MCI. These findings suggest that PET and CSF do not provide identical information about the stage of Alzheimer’s disease.

The idea to study differences in the predictive strength of patient features for PET/CSF amyloid-β status was based on the differences in characteristics of patients with discordant amyloid-β biomarkers, which have been theorized to be caused by various factors. Possible explanations for the discordance include individual variances in CSF Aβ_42_ production (55), the composition of amyloid-β plaques (56), differences in the structure of Aβ fibrils (57), or a variety of technical issues (58,59), including the variability in cut-off values for CSF Aβ_42_ (14). It has also been proposed that in the earliest stages of amyloid-β accumulation CSF Aβ_42_ analysis might be more sensitive, as the decrease in the concentration of soluble isoforms might precede fibrillar amyloid-β plaque deposition detectable by PET (21). The differences in the predictive pattern found here in relation to other biological variables, such as APOE genotype and (p)tau concentrations, imply that the existence of PET/CSF discordance may not only be due to technical variation, but a reflection of the differences in biological substrate between the modalities. This could also have an effect for future practice in AD research as well as patient care, as the two modalities are currently used as equal alternatives (2,11).

We observed that CSF p-tau and tau had a stronger association to amyloid-β based on PET compared to CSF. If we assume that CSF is a more sensitive modality for amyloid-β pathology, then the weaker association with tau could be explained by CSF Aβ_42_ capturing an earlier stage amyloid-β preceding tau pathology. This was reflected by the predictive patterns in the multivariable logistic regression models: when predicting PET status by CSF status, CSF (p)tau adds information about the added burden of disease (including advancing from CSF+PET- to CSF+PET+). When predicting CSF amyloid-β positivity, however, the existence of amyloid-β pathology on PET already provides sufficient predictive power, of subjects already having reached a later stage in amyloid deposition. Overall, this finding is in accordance with previous work, indicating that PET detects more advanced stages of AD pathology (60). Although CSF tau and p-tau have been shown to be highly correlated (61), the results of the random forest models imply that CSF tau is more predictive towards amyloid-β pathology in SCD and MCI, whereas CSF p-tau is more predictive in dementia. This finding might be caused by wider neuronal death preceding the release of phosphorylated tau, although previous work seems to suggest that levels of CSF p-tau decrease in the later stages of AD (62–64). Another possible explanation is that this finding is caused by the greater specificity of p-tau for AD pathology (65), as our cohort also included amyloid-positive patients diagnosed with non-AD dementia, possibly due to secondary amyloid pathology.

Although we focus on the relative differences between PET and CSF, it should be emphasized that in the majority of cases these two modalities contain similar information. This was demonstrated by many of the selected patient features having some predictive power for amyloid-β pathology for both PET and CSF. Of them, the biological factors *APOE* ε4 carriership, CSF tau, and p-tau were most consistent in having significant predictive ability amyloid-β status irrespective of the modality. These findings are not unexpected, as *APOE* ε4 carriership (17,66,67) and tau pathology (2,68) are widely known to have a strong connection to amyloid-β pathology in Alzheimer’s disease. Cognitive measures and MRI visual reads showed overall a smaller predictive value towards amyloid-β status, being in concordance with the theory that they show changes downstream of amyloid and tau pathology (69).

The main strength of our study is the large number of patients with both amyloid-β modalities from a well-characterized cohort. Our study also has several weaknesses. First, due to the stratification by syndrome diagnoses, the outcome of amyloid-β positivity was not equally prevalent. Although we used an AUC-based importance measure that has been shown to be more effective with an unbalanced outcome class (52), there was still a notably low sensitivity in the SCD group. This could theoretically influence the outcome of the random forest models, although we found comparable results when using logistic regression models. Second, the included patients underwent amyloid-β PET scans with four different radiotracers, allowing for variability in thresholds for amyloid-β positivity. However, this effect is likely reduced by all of the PET scans being visually rated by the same experienced nuclear medicine physician. Third, as continuous measures for PET-imaging were not available, we dichotomized CSF Aβ_42_ values, causing some loss of information. Fourth, this patient group did not have CSF Aβ_40_ values available, which have been shown to correct for the individual variation in the production of amyloid-β (70,71).

Our findings can be summarized by a hypothetical model highlighting the relative predictive power of patient features towards amyloid-β status based on PET and CSF (**Figure 3**). This model supports previous work, suggesting that CSF might be more sensitive in the early stages of amyloid-β pathology, whereas PET status might be more specific to later stages of amyloid-β accumulation. Although the modalities show similar information in the majority of cases, this could have implications for future research and clinical trials. For example, if aiming to capture the earliest stage of amyloid-β pathology, CSF might be preferred over PET. On the contrary, if high confidence of significant amyloid-β pathology is required, PET could be the modality of choice.

**Figure 3.**
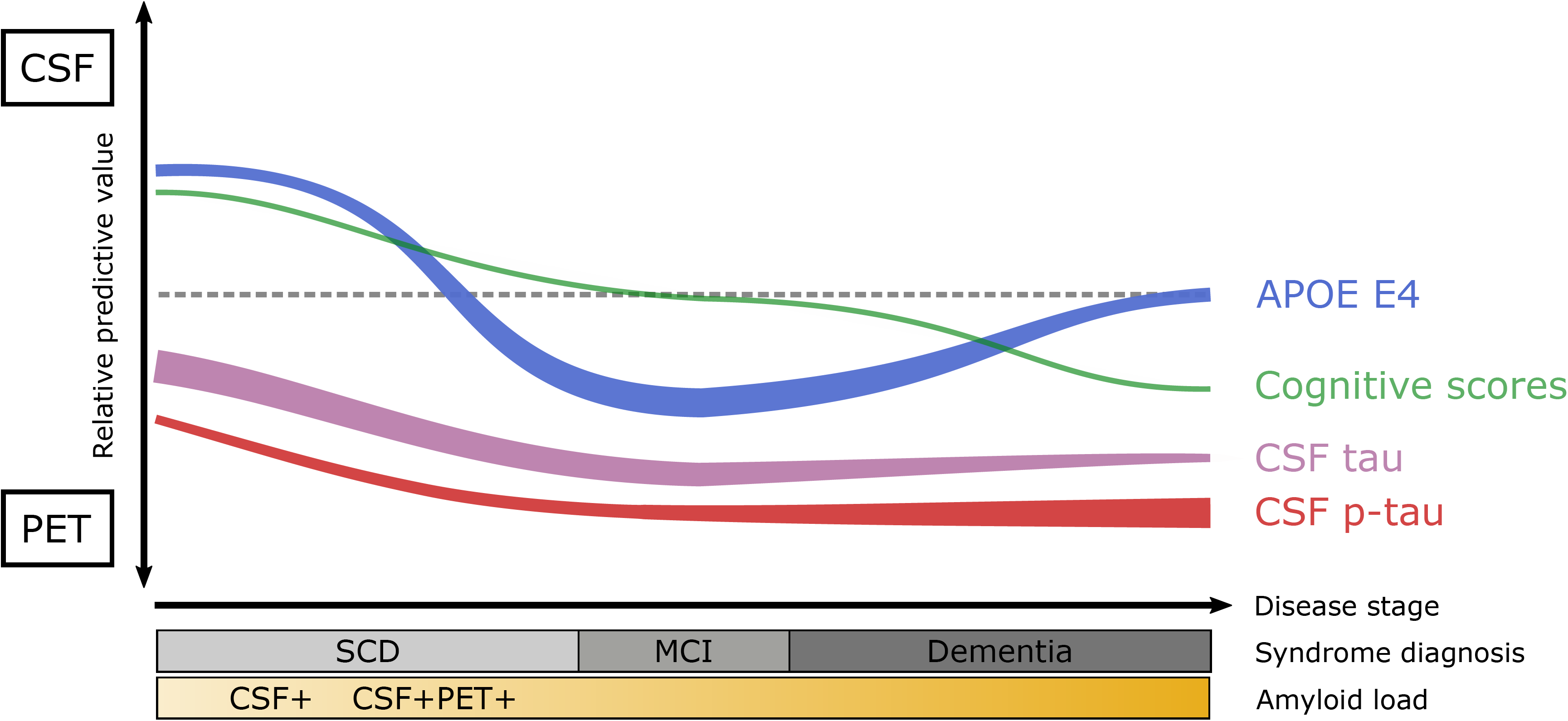
Hypothetical model for relative predictive strength of patient features toward PET and CSF amyloid status. Line location on the y-axis indicates the relative strength of the association between the patient feature and status of the amyloid-□ modality. Line thickness indicates the overall predictive strength of the patient feature for amyloid status based on both PET and CSF.

## CONCLUSION

We demonstrated that although various patient features have general predictive value towards amyloid-β status, there are finer differences revealed by discordant cases between the predictive pattern for amyloid-β status based on PET and CSF. This indicates that PET-CSF discordance includes valuable information on underlying clinical and neuropathological differences.

## DECLARATIONS

The local medical ethics committee of VU University Medical Center has approved a general protocol for biobanking and using the clinical data for research purposes. The data used in this study are not publicly available but may be provided upon reasonable request. The Alzheimer Center Amsterdam is supported by Alzheimer Nederland and Stichting VUmc fonds. Research performed at the Alzheimer Center Amsterdam is part of the neurodegeneration research program of Amsterdam Neuroscience. The clinical database structure was developed with funding from Stichting Dioraphte. WvdfF was recipient of ZonMW-Memorabel ABIDE (project No 733050201), a project in the context of the Dutch Deltaplan Dementie. WvdF holds the Pasman chair. The funding sources had no involvement in the writing of this article or in the decision to submit it for publications. JR acknowledges Prof Sergei Nazarenko, the North Estonia Medical Centre and the International Atomic Energy Agency for their contribution to his professional development.

## AUTHORS’ CONTRIBUTIONS

JR, RO, and FBo conceived the study, designed the protocol, analyzed/interpreted data, drafted the manuscript. JR performed statistical analysis. RO, FBo, and PS provided overall study supervision. AdW, CET, MZ, ADW, RB, FBa, WMvdF, PS, BNMvB had a major role in the acquisition of data, and critically revised and edited the manuscript for intellectual content. All authors read and approved the final version of the manuscript.

