## Supplemental Tables for "PET and CSF amyloid-β status are differently predicted by patient features: Information from discordant cases"

**Supplementary Table 1**. Proportion of missing values per patient feature

| | PET-CSF- | PET+CSF- | PET-CSF+ | PET+CSF+ |
|---|---|---|---|---|
| n | 315 | 32 | 65 | 356 |
| Age (%) | 0 (0.0) | 0 (0.0) | 0 (0.0) | 0 (0.0) |
| Sex (%) | 0 (0.0) | 0 (0.0) | 0 (0.0) | 0 (0.0) |
| Education (%) | 13 (4.1) | 2 (6.2) | 3 (4.6) | 10 (2.8) |
| APOE ε4 (%) | 12 (3.8) | 2 (6.2) | 4 (6.2) | 14 (3.9) |
| CSF tau (%) | 0 (0.0) | 0 (0.0) | 1 (1.5) | 2 (0.6) |
| CSF p-tau (%) | 2 (0.6) | 0 (0.0) | 4 (6.2) | 0 (0.0) |
| MMSE (%) | 5 (1.6) | 1 (3.1) | 2 (3.1) | 7 (2.0) |
| Memory z-score (%) | 14 (4.4) | 3 (9.4) | 3 (4.6) | 21 (5.9) |
| Language z-score (%) | 17 (5.4) | 3 (9.4) | 3 (4.6) | 25 (7.0) |
| Attention z-score (%) | 15 (4.8) | 3 (9.4) | 3 (4.6) | 22 (6.2) |
| Executive z-score (%) | 6 (1.9) | 1 (3.1) | 2 (3.1) | 12 (3.4) |
| Visuospatial z-score (%) | 23 (7.3) | 3 (9.4) | 5 (7.7) | 36 (10.1) |
| MRI MTA (%) | 68 (21.6) | 4 (12.5) | 6 (9.2) | 88 (24.7) |
| MRI PCA (%) | 91 (28.9) | 4 (12.5) | 7 (10.8) | 93 (26.1) |
| MRI Fazekas (%) | 68 (21.6) | 4 (12.5) | 2 (3.1) | 89 (25.0) |
| MRI lacunes (%) | 76 (24.1) | 5 (15.6) | 3 (4.6) | 95 (26.7) |
| MRI microbleeds (%) | 78 (24.8) | 5 (15.6) | 10 (15.4) | 102 (28.7) |

**Supplementary Table 2**. Out-of-bag accuracy, sensitivity and specificity for random forest models predicting amyloid PET and CSF status

| | Outcome | Accuracy % | Sensitivity % | Specificity % |
|---|---|---|---|---|
| **Total** | PET | 82 (81, 83) | 82 (81, 83) | 82 (81, 83) |
| | CSF | 78 (77, 78) | 81 (80, 82) | 74 (73, 75) |
| **SCD** | PET | 82 (80, 82) | 22 (18, 26) | 96 (96, 96) |
| | CSF | 79 (77, 80) | 31 (27, 36) | 94 (94, 95) |
| **MCI** | PET | 82 (80, 84) | 81 (79, 83) | 84 (82, 84) |
| | CSF | 72 (69, 75) | 74 (70, 77) | 70 (65, 75) |
| **Dementia** | PET | 82 (82, 83) | 90 (89, 91) | 69 (68, 71) |
| | CSF | 78 (77, 80) | 90 (89, 91) | 53 (50, 57) |

Mean rates with 95% confidence intervals over 25 random forest models are reported.

**Supplementary Table 3.** Predictive value of patient features for amyloid status based on PET or CSF

| | | TOTAL | | | | | SCD | | | | | MCI | | | | | DEMENTIA | | | | |
|---|---|---|---|---|---|---|---|---|---|---|---|---|---|---|---|---|---|---|---|---|---|
| | | | | | Imputed | | | | | Imputed | | | | | Imputed | | | | | Imputed | |
| **Predictor** | **Out-come** | **Odds ratio (95% CI)** | **p unc** | **p FDR** | **Odds ratio (95% CI)** | **p FDR** | **Odds ratio (95% CI)** | **p unc** | **p FDR** | **Odds ratio (95% CI)** | **p FDR** | **Odds ratio (95% CI)** | **p unc** | **p FDR** | **Odds ratio (95% CI)** | **p FDR** | **Odds ratio (95% CI)** | **p unc** | **p FDR** | **Odds ratio (95% CI)** | **p FDR** |
| Age | PET | 1.02(1.00,1.04) | | | 1.02(1.00,1.04) | | 1.06(1.01,1.12) | * | | 1.06(1.01,1.12) | | 0.96(0.92,1.00) | | | 0.96(0.92,1.00) | | 1.00(0.97,1.03) | | | 1.00(0.97,1.03) | |
| | CSF | 1.01(0.99,1.03) | | | 1.01(0.99,1.03) | | 1.04(1.00,1.09) | | | 1.04(1.00,1.09) | | 0.96(0.92,1.00) | | | 0.96(0.92,1.00) | | 0.98(0.96,1.01) | | | 0.98(0.96,1.01) | |
| Sex, F | PET | 1.69(1.26,2.26) | *** | *** | 1.69(1.26,2.26) | *** | 1.64(0.81,3.36) | | | 1.64(0.81,3.36) | | 2.45(1.14,5.26) | * | | 2.45(1.14,5.26) | | 1.70(1.13,2.53) | ** | * | 1.70(1.13,2.53) | * |
| | CSF | 1.55(1.15,2.07) | ** | ** | 1.55(1.15,2.07) | ** | 1.91(0.99,3.69) | | | 1.91(0.99,3.69) | | 2.01(0.94,4.28) | | | 2.01(0.94,4.28) | | 1.40(0.93,2.12) | | | 1.40(0.93,2.12) | |
| Education | PET | 1.06(0.94,1.19) | | | 1.07(0.95,1.20) | | 1.10(0.84,1.46) | | | 1.10(0.84,1.45) | | 0.92(0.69,1.24) | | | 0.91(0.68,1.22) | | 1.28(1.08,1.52) | ** | ** | 1.30(1.10,1.53) | ** |
| | CSF | 1.04(0.93,1.17) | | | 1.05(0.94,1.18) | | 1.05(0.82,1.35) | | | 1.04(0.81,1.34) | | 0.93(0.69,1.24) | | | 0.94(0.71,1.26) | | 1.29(1.08,1.54) | ** | * | 1.30(1.09,1.54) | ** |
| APOE E4 | PET | 4.72(3.46,6.44) | *** | *** | 4.57(3.34,6.24) | *** | 2.97(1.42,6.20) | ** | * | 2.97(1.42,6.19) | * | 14.55(6.08,34.82) | *** | *** | 13.43(5.62,32.11) | *** | 3.63(2.39,5.50) | *** | *** | 3.51(2.30,5.34) | *** |
| | CSF | 4.60(3.36,6.28) | *** | *** | 4.46(3.26,6.12) | *** | 3.82(1.90,7.70) | *** | ** | 3.75(1.86,7.57) | ** | 8.28(3.70,18.54) | *** | *** | 7.76(3.49,17.27) | *** | 3.68(2.39,5.69) | *** | *** | 3.59(2.33,5.53) | *** |
| CSF tau | PET | 1.005 (1.004,1.006) | *** | *** | 1.005 (1.004,1.006) | *** | 1.004 (1.002,1.006) | *** | *** | 1.004 (1.002,1.006) | *** | 1.008 (1.005,1.011) | *** | *** | 1.008 (1.005,1.011) | *** | 1.004 (1.003,1.005) | *** | *** | 1.004 (1.003,1.005) | *** |
| | CSF | 1.004 (1.003,1.005) | *** | *** | 1.004 (1.003,1.005) | *** | 1.003 (1.002,1.005) | *** | ** | 1.003 (1.002,1.005) | ** | 1.003 (1.002,1.005) | *** | *** | 1.003 (1.002,1.005) | ** | 1.004 (1.003,1.004) | *** | *** | 1.003 (1.002,1.004) | *** |
| CSF p-tau | PET | 1.05(1.04,1.06) | *** | *** | 1.05(1.04,1.06) | *** | 1.04(1.02,1.05) | *** | *** | 1.04(1.02,1.05) | *** | 1.05(1.03,1.07) | *** | *** | 1.05(1.03,1.07) | *** | 1.05(1.04,1.06) | *** | *** | 1.05(1.04,1.06) | *** |
| | CSF | 1.04(1.03,1.04) | *** | *** | 1.04(1.03,1.04) | *** | 1.03(1.01,1.04) | *** | ** | 1.02(1.01,1.04) | ** | 1.03(1.01,1.04) | *** | *** | 1.03(1.01,1.04) | ** | 1.04(1.03,1.05) | *** | *** | 1.04(1.03,1.05) | *** |
| MMSE | PET | 1.20(1.15,1.25) | *** | *** | 1.19(1.14,1.24) | *** | 1.03(0.89,1.19) | | | 1.02(0.88,1.18) | | 1.11(0.95,1.30) | | | 1.12(0.95,1.31) | | 1.12(1.06,1.18) | *** | *** | 1.12(1.06,1.18) | *** |
| | CSF | 1.20(1.15,1.25) | *** | *** | 1.19(1.14,1.25) | *** | 1.15(1.01,1.31) | * | | 1.13(1.00,1.29) | | 1.02(0.87,1.20) | | | 1.02(0.87,1.19) | | 1.10(1.04,1.17) | *** | ** | 1.10(1.04,1.17) | ** |
| Memory | PET | 1.36(1.27,1.47) | *** | *** | 1.36(1.27,1.46) | *** | 1.13(0.82,1.55) | | | 1.14(0.83,1.56) | | 1.26(1.02,1.57) | * | | 1.25(1.00,1.54) | | 1.20(1.10,1.30) | *** | *** | 1.20(1.10,1.31) | *** |
| | CSF | 1.32(1.23,1.42) | *** | *** | 1.32(1.23,1.42) | *** | 1.23(0.92,1.64) | | | 1.22(0.91,1.62) | | 1.16(0.95,1.42) | | | 1.12(0.92,1.38) | | 1.14(1.05,1.24) | ** | ** | 1.15(1.06,1.26) | ** |
| Language | PET | 1.10(1.00,1.20) | | | 1.09(0.99,1.20) | | 0.95(0.56,1.60) | | | 0.94(0.56,1.57) | | 0.38(0.18,0.81) | * | | 0.44(0.21,0.95) | | 0.94(0.85,1.04) | | | 0.94(0.85,1.04) | |
| | CSF | 1.20(1.07,1.34) | ** | ** | 1.19(1.07,1.33) | ** | 0.99(0.62,1.58) | | | 0.99(0.62,1.56) | | 0.71(0.39,1.27) | | | 0.71(0.39,1.29) | | 1.00(0.90,1.11) | | | 1.00(0.90,1.11) | |
| Attention | PET | 1.31(1.14,1.49) | *** | *** | 1.27(1.12,1.45) | *** | 1.06(0.72,1.55) | | | 1.00(0.69,1.46) | | 0.57(0.34,0.94) | * | | 0.66(0.41,1.05) | | 1.03(0.86,1.23) | | | 1.01(0.85,1.19) | |
| | CSF | 1.36(1.19,1.56) | *** | *** | 1.32(1.16,1.50) | *** | 1.10(0.77,1.55) | | | 1.07(0.76,1.50) | | 0.86(0.54,1.37) | | | 0.98(0.62,1.53) | | 1.00(0.83,1.20) | | | 0.97(0.81,1.16) | |
| Executive | PET | 1.28(1.15,1.42) | *** | *** | 1.28(1.15,1.42) | *** | 1.02(0.72,1.45) | | | 1.02(0.72,1.44) | | 0.68(0.45,1.04) | | | 0.70(0.46,1.05) | | 0.95(0.82,1.11) | | | 0.96(0.82,1.11) | |
| | CSF | 1.27(1.14,1.41) | *** | *** | 1.27(1.14,1.42) | *** | 1.05(0.76,1.44) | | | 1.05(0.76,1.44) | | 0.82(0.55,1.22) | | | 0.83(0.55,1.24) | | 0.88(0.76,1.03) | | | 0.89(0.76,1.04) | |
| Visuo-spatial | PET | 1.36(1.22,1.52) | *** | *** | 1.33(1.19,1.48) | *** | 0.92(0.59,1.45) | | | 0.89(0.56,1.41) | | 0.72(0.47,1.08) | | | 0.78(0.52,1.16) | | 1.30(1.15,1.49) | *** | *** | 1.25(1.10,1.43) | ** |
| | CSF | 1.38(1.23,1.55) | *** | *** | 1.34(1.19,1.50) | *** | 1.34(0.93,1.93) | | | 1.35(0.95,1.93) | | 0.98(0.68,1.42) | | | 1.02(0.71,1.47) | | 1.21(1.07,1.37) | ** | ** | 1.16(1.03,1.31) | * |
| MRI MTA | PET | 1.27(1.05,1.53) | * | * | 1.25(1.04,1.50) | * | 1.78(0.90,3.53) | | | 1.56(0.80,3.04) | | 0.77(0.48,1.24) | | | 0.75(0.47,1.20) | | 0.77(0.60,1.00) | | | 0.81(0.63,1.04) | |
| | CSF | 1.40(1.16,1.70) | *** | ** | 1.34(1.11,1.61) | ** | 1.47(0.76,2.84) | | | 1.37(0.72,2.59) | | 1.09(0.69,1.73) | | | 0.98(0.62,1.54) | | 0.81(0.62,1.05) | | | 0.84(0.65,1.09) | |
| MRI PCA | PET | 1.65(1.32,2.07) | *** | *** | 1.62(1.30,2.01) | *** | 1.71(0.93,3.16) | | | 1.55(0.87,2.78) | | 0.84(0.47,1.51) | | | 0.85(0.48,1.51) | | 1.20(0.88,1.62) | | | 1.18(0.87,1.59) | |
| | CSF | 1.38(1.11,1.73) | ** | ** | 1.41(1.14,1.74) | ** | 1.07(0.60,1.90) | | | 1.08(0.61,1.90) | | 0.74(0.41,1.32) | | | 0.75(0.42,1.32) | | 0.97(0.70,1.34) | | | 1.02(0.75,1.40) | |
| MRI Fazekas | PET | 1.02(0.82,1.27) | | | 1.01(0.81,1.25) | | 0.98(0.53,1.83) | | | 0.92(0.50,1.71) | | 0.83(0.48,1.42) | | | 0.78(0.45,1.38) | | 0.80(0.60,1.06) | | | 0.83(0.62,1.11) | |
| | CSF | 1.19(0.95,1.48) | | | 1.12(0.90,1.39) | | 1.38(0.79,2.41) | | | 1.25(0.72,2.16) | | 0.77(0.45,1.32) | | | 0.74(0.42,1.30) | | 0.96(0.71,1.31) | | | 0.95(0.70,1.30) | |
| MRI Lacunes | PET | 0.84(0.43,1.62) | | | 0.85(0.44,1.65) | | 0(0,Inf) | | | 0(0,Inf) | | 0.33(0.06,1.70) | | | 0.36(0.07,1.83) | | 0.72(0.33,1.59) | | | 0.73(0.33,1.63) | |
| | CSF | 1.44(0.73,2.85) | | | 1.25(0.63,2.46) | | 6.16(0.54,69.92) | | | 3.58(0.35,36.7) | | 0.31(0.06,1.63) | | | 0.37(0.07,2.00) | | 1.25(0.51,3.05) | | | 1.06(0.44,2.58) | |
| MRI microbleeds | PET | 1.91(1.21,3.01) | ** | ** | 1.59(1.01,2.50) | | 2.17(0.75,6.30) | | | 1.84(0.68,4.99) | | 1.32(0.52,3.32) | | | 1.11(0.47,2.62) | | 2.36(1.16,4.81) | * | * | 1.93(0.93,3.99) | |
| | CSF | 1.51(0.96,2.39) | | | 1.35(0.86,2.10) | | 1.49(0.52,4.25) | | | 1.32(0.49,3.52) | | 1.06(0.42,2.65) | | | 0.98(0.41,2.32) | | 1.98(0.95,4.13) | | | 1.70(0.81,3.56) | |

*** - p<0.001, ** - p<0.01, * - p<0.05. P-values indicate the significance of the patient feature in the model. Uncorrected p-values and corrected p-values are reported per model, additionally corrected p-values for imputed data. False discovery rate (FDR) correction was performed for multiple comparisons. Cognitive scores have been multiplied by -1, therefore lower scores usually indicate higher odds ratios for amyloid positivity.

**Supplementary Table 4**. Amyloid-adjusted predictive value of patient features for amyloid status based on PET or CSF

| | | TOTAL | | | | | SCD | | | | | MCI | | | | | DEMENTIA | | | | |
|---|---|---|---|---|---|---|---|---|---|---|---|---|---|---|---|---|---|---|---|---|---|
| | | | | | Imputed | | | | | Imputed | | | | | Imputed | | | | | Imputed | |
| Predictor | Out-come | Odds ratio (95% CI) | p unc | p FDR | Odds ratio (95% CI) | p FDR | Odds ratio (95% CI) | p unc | p FDR | Odds ratio (95% CI) | p FDR | Odds ratio (95% CI) | p unc | p FDR | Odds ratio (95% CI) | p FDR | Odds ratio (95% CI) | p unc | p FDR | Odds ratio (95% CI) | p FDR |
| Age | PET | 1.03(1.00,1.06) | | | 1.03(1.00,1.06) | | 1.04(0.99,1.10) | | | 1.04(0.99,1.10) | | 0.97(0.91,1.04) | | | 0.97(0.91,1.04) | | 1.03(0.99,1.07) | | | 1.03(0.99,1.07) | |
| | CSF | 0.99(0.96,1.02) | | | 0.99(0.96,1.02) | | 1.02(0.97,1.07) | | | 1.02(0.97,1.07) | | 0.98(0.92,1.04) | | | 0.98(0.92,1.04) | | 0.96(0.93,1.00) | | | 0.96(0.93,1.00) | |
| Sex, F | PET | 1.57(1.01,2.44) | * | | 1.57(1.01,2.44) | | 1.17(0.49,2.77) | | | 1.17(0.49,2.77) | | 2.27(0.75,6.90) | | | 2.27(0.75,6.90) | | 1.93(1.04,3.58) | * | | 1.93(1.04,3.58) | |
| | CSF | 1.10(0.71,1.72) | | | 1.10(0.71,1.72) | | 1.76(0.80,3.90) | | | 1.76(0.80,3.90) | | 1.11(0.37,3.35) | | | 1.11(0.37,3.35) | | 0.84(0.44,1.59) | | | 0.84(0.44,1.59) | |
| Education | PET | 1.06(0.89,1.27) | | | 1.07(0.89,1.27) | | 1.11(0.79,1.56) | | | 1.12(0.79,1.57) | | 0.95(0.60,1.50) | | | 0.90(0.57,1.41) | | 1.16(0.90,1.49) | | | 1.18(0.92,1.51) | |
| | CSF | 1.00(0.84,1.19) | | | 1.00(0.84,1.19) | | 1.00(0.74,1.35) | | | 0.99(0.73,1.34) | | 0.97(0.61,1.54) | | | 1.02(0.65,1.60) | | 1.15(0.89,1.50) | | | 1.14(0.88,1.48) | |
| APOE E4 | PET | 2.58(1.65,4.03) | *** | *** | 2.52(1.62,3.93) | *** | 1.54(0.62,3.78) | | | 1.56(0.63,3.82) | | 9.44(2.93,30.39) | *** | ** | 8.79(2.72,28.41) | ** | 2.22(1.20,4.09) | * | | 2.14(1.16,3.95) | |
| | CSF | 2.30(1.47,3.60) | *** | ** | 2.28(1.45,3.57) | ** | 3.07(1.33,7.07) | ** | | 3.01(1.30,6.94) | | 1.85(0.58,5.92) | | | 1.85(0.58,5.88) | | 2.00(1.06,3.78) | * | | 2.00(1.07,3.75) | |
| CSF tau | PET | 1.003 (1.003,1.004) | *** | *** | 1.003 (1.003,1.004) | *** | 1.003 (1.001,1.005) | ** | * | 1.003 (1.001,1.005) | * | 1.008 (1.004,1.012) | *** | ** | 1.008 (1.004,1.012) | ** | 1.003 (1.002,1.004) | *** | *** | 1.003 (1.002,1.004) | *** |
| | CSF | 1.001 (1.000,1.002) | ** | * | 1.001 (1.000,1.002) | * | 1.002 (1.000,1.003) | | | 1.001 (1.000,1.003) | | 0.999 (0.997,1.001) | | | 0.999 (0.997,1.001) | | 1.001 (1.000,1.002) | | | 1.001 (1.000,1.002) | |
| CSF p-tau | PET | 1.04(1.03,1.05) | *** | *** | 1.04(1.03,1.05) | *** | 1.02(1.01,1.04) | ** | * | 1.03(1.01,1.04) | * | 1.05(1.02,1.07) | *** | ** | 1.05(1.02,1.07) | ** | 1.04(1.03,1.05) | *** | *** | 1.04(1.03,1.05) | *** |
| | CSF | 1.01(1.00,1.02) | * | | 1.01(1.00,1.02) | | 1.01(1.00,1.03) | | | 1.01(0.99,1.02) | | 0.99(0.98,1.01) | | | 0.99(0.98,1.01) | | 1.01(1.00,1.02) | | | 1.01(1.00,1.02) | |
| MMSE | PET | 1.11(1.05,1.17) | *** | ** | 1.10(1.04,1.17) | ** | 0.93(0.80,1.10) | | | 0.93(0.79,1.09) | | 1.22(0.96,1.56) | | | 1.24(0.97,1.59) | | 1.10(1.02,1.19) | * | | 1.10(1.01,1.18) | |
| | CSF | 1.10(1.04,1.16) | ** | ** | 1.10(1.04,1.16) | ** | 1.21(1.03,1.41) | * | | 1.19(1.02,1.38) | | 0.88(0.69,1.12) | | | 0.87(0.69,1.11) | | 1.02(0.94,1.10) | | | 1.02(0.94,1.11) | |
| Memory | PET | 1.22(1.12,1.34) | *** | *** | 1.22(1.12,1.33) | *** | 0.99(0.69,1.42) | | | 1.01(0.70,1.46) | | 1.25(0.96,1.64) | | | 1.27(0.97,1.65) | | 1.18(1.05,1.32) | ** | * | 1.17(1.05,1.31) | |
| | CSF | 1.09(1.00,1.19) | * | | 1.09(1.01,1.19) | | 1.23(0.87,1.75) | | | 1.21(0.85,1.72) | | 0.96(0.71,1.30) | | | 0.92(0.68,1.25) | | 1.00(0.89,1.11) | | | 1.01(0.91,1.12) | |
| Language | PET | 0.95(0.85,1.07) | | | 0.95(0.84,1.07) | | 0.91(0.45,1.86) | | | 0.91(0.46,1.80) | | 0.23(0.08,0.68) | ** | | 0.32(0.10,1.01) | | 0.90(0.79,1.01) | | | 0.89(0.79,1.01) | |
| | CSF | 1.24(1.08,1.43) | ** | * | 1.23(1.07,1.42) | * | 1.03(0.58,1.82) | | | 1.02(0.59,1.78) | | 1.59(0.77,3.27) | | | 1.37(0.63,2.98) | | 1.12(0.95,1.32) | | | 1.13(0.96,1.34) | |
| Attention | PET | 1.10(0.91,1.34) | | | 1.09(0.90,1.32) | | 1.00(0.65,1.52) | | | 0.96(0.63,1.45) | | 0.38(0.18,0.80) | * | | 0.43(0.21,0.86) | | 1.07(0.81,1.40) | | | 1.07(0.82,1.39) | |
| | CSF | 1.27(1.03,1.55) | * | | 1.24(1.02,1.50) | | 1.10(0.71,1.70) | | | 1.09(0.72,1.66) | | 1.80(0.88,3.68) | | | 1.87(0.93,3.77) | | 0.95(0.72,1.26) | | | 0.92(0.70,1.21) | |
| Executive | PET | 1.17(1.00,1.37) | | | 1.16(1.00,1.36) | | 0.99(0.67,1.48) | | | 0.99(0.67,1.47) | | 0.61(0.33,1.12) | | | 0.62(0.34,1.14) | | 1.11(0.88,1.40) | | | 1.10(0.87,1.39) | |
| | CSF | 1.12(0.96,1.31) | | | 1.13(0.97,1.32) | | 1.05(0.71,1.55) | | | 1.05(0.72,1.54) | | 1.18(0.64,2.17) | | | 1.17(0.64,2.15) | | 0.81(0.64,1.03) | | | 0.82(0.64,1.04) | |
| Visuo-spatial | PET | 1.19(1.03,1.37) | * | * | 1.17(1.03,1.34) | | 0.77(0.49,1.22) | | | 0.73(0.46,1.18) | | 0.55(0.31,0.96) | * | | 0.61(0.36,1.03) | | 1.32(1.10,1.59) | ** | * | 1.28(1.07,1.53) | |
| | CSF | 1.20(1.04,1.39) | * | * | 1.16(1.01,1.34) | | 1.53(0.99,2.38) | | | 1.58(1.01,2.45) | | 1.63(0.93,2.83) | | | 1.53(0.89,2.63) | | 0.99(0.84,1.17) | | | 0.97(0.83,1.13) | |
| MRI MTA | PET | 1.00(0.76,1.31) | | | 1.02(0.78,1.33) | | 1.68(0.77,3.68) | | | 1.48(0.66,3.32) | | 0.55(0.28,1.05) | | | 0.58(0.30,1.09) | | 0.79(0.55,1.15) | | | 0.81(0.56,1.17) | |
| | CSF | 1.37(1.06,1.77) | * | * | 1.30(1.01,1.68) | | 1.15(0.53,2.51) | | | 1.12(0.51,2.43) | | 1.78(0.87,3.63) | | | 1.52(0.76,3.03) | | 0.96(0.67,1.38) | | | 0.99(0.69,1.41) | |
| MRI PCA | PET | 1.71(1.25,2.33) | *** | ** | 1.66(1.22,2.26) | ** | 1.87(0.93,3.76) | | | 1.74(0.88,3.43) | | 1.09(0.48,2.46) | | | 1.11(0.50,2.48) | | 1.50(0.97,2.31) | | | 1.42(0.93,2.17) | |
| | CSF | 0.95(0.70,1.29) | | | 0.96(0.72,1.30) | | 0.78(0.39,1.56) | | | 0.78(0.39,1.57) | | 0.69(0.30,1.58) | | | 0.69(0.31,1.52) | | 0.74(0.47,1.14) | | | 0.79(0.51,1.21) | |
| MRI Fazekas | PET | 0.82(0.61,1.11) | | | 0.84(0.62,1.14) | | 0.76(0.36,1.62) | | | 0.70(0.33,1.50) | | 0.98(0.46,2.08) | | | 0.95(0.45,2.02) | | 0.68(0.46,1.00) | | | 0.72(0.48,1.07) | |
| | CSF | 1.35(1.00,1.81) | | | 1.26(0.94,1.70) | | 1.56(0.83,2.96) | | | 1.48(0.78,2.78) | | 0.78(0.37,1.64) | | | 0.76(0.36,1.61) | | 1.28(0.85,1.94) | | | 1.23(0.81,1.87) | |
| MRI Lacunes | PET | 0.47(0.20,1.09) | | | 0.51(0.22,1.20) | | 0(0,Inf) | | | 0(0,Inf) | | 0.54(0.06,4.90) | | | 0.53(0.07,4.09) | | 0.44(0.16,1.20) | | | 0.47(0.17,1.33) | |
| | CSF | 2.42(1.01,5.78) | * | | 2.03(0.85,4.86) | | 12.35(1.06,143.8) | * | | 7.61(0.69,84.47) | | 0.47(0.05,4.24) | | | 0.58(0.07,4.96) | | 2.28(0.73,7.12) | | | 1.89(0.60,5.95) | |
| MRI microbleeds | PET | 2.08(1.07,4.03) | * | | 1.75(0.90,3.41) | | 2.15(0.60,7.67) | | | 1.94(0.55,6.86) | | 1.62(0.43,6.12) | | | 1.30(0.36,4.68) | | 2.24(0.82,6.12) | | | 1.87(0.69,5.09) | |
| | CSF | 0.89(0.46,1.73) | | | 0.88(0.46,1.71) | | 1.01(0.29,3.57) | | | 0.91(0.26,3.17) | | 0.75(0.20,2.81) | | | 0.81(0.22,2.94) | | 1.08(0.38,3.09) | | | 1.04(0.38,2.88) | |

*** - p<0.001, ** - p<0.01, * - p<0.05. P-values indicate the significance of the patient feature in the model. Uncorrected p-values and corrected p-values are reported per model, additionally corrected p-values for imputed data. False discovery rate (FDR) correction was performed for multiple comparisons. Cognitive scores have been multiplied by -1, therefore lower scores usually indicate higher odds ratios for amyloid positivity.
